# Mechanism of ribosome rescue by ArfA and RF2

**DOI:** 10.1101/091256

**Authors:** Gabriel Demo, Egor Svidritskiy, Rohini Madireddy, Ruben Diaz-Avalos, Timothy Grant, Nikolaus Grigorieff, Duncan Sousa, Andrei A. Korostelev

**Affiliations:** RNA Therapeutics Institute, University of Massachusetts Medical School, 368 Plantation St., Worcester, MA 01605, USA; Department of Biochemistry and Molecular Pharmacology, University of Massachusetts Medical School, 368 Plantation St., Worcester, MA 01605, USA; Janelia Research Campus, Howard Hughes Medical Institute, 19700 Helix Drive, Ashburn, VA 20147, USA; Department of Biological Science, Florida State University, 89 Chieftan Way, Tallahassee, FL 32306, USA

**Author notes:** Current address: Medicago, 7 Triangle Dr, Durham, NC 27707. **Materials & Correspondence**: Andrei A. Korostelev, University of Massachusetts Medical School, 368 Plantation Street, Worcester MA 01605, USA.

## Abstract

ArfA rescues ribosomes stalled on truncated mRNAs by recruiting the release factor RF2, which normally binds stop codons to catalyze peptide release. We report two 3.2-Å resolution cryo-EM structures – determined from a single sample – of the 70S ribosome with ArfA∙RF2 in the A site. In both states, the ArfA C-terminus occupies the mRNA tunnel downstream of the A site. One state contains a compact inactive RF2 conformation, hitherto unobserved in 70S termination complexes. Ordering of the ArfA N-terminus in the second state rearranges RF2 into an extended conformation that docks the catalytic GGQ motif into the peptidyl-transferase center. Our work thus reveals the structural dynamics of ribosome rescue. The structures demonstrate how ArfA “senses” the vacant mRNA tunnel and activates RF2 to mediate peptide release without a stop codon, allowing stalled ribosomes to be recycled.

## Introduction

A translating ribosome stalls when it encounters the end of a non-stop mRNA, truncated during cellular stress or homeostasis, by premature transcription termination or mRNA cleavage or other mechanisms (Hayes and Keiler 2010; Keiler 2015). The stalled ribosome contains *<u>p</u>*eptidyl-tRNA in the *P* site, whereas the *<u>a</u>*minoacyl-tRNA (*A*) site is unoccupied (Ito et al. 2011). Bacteria have evolved ribosome-rescue pathways to release the nascent peptide and re-enable the stalled ribosome for translation (for a review, see ref. (Keiler 2015)). The ArfA (<u>a</u>lternative <u>r</u>escue <u>f</u>actor <u>A</u>) pathway is essential in trans-translation-deficient cells (Chadani et al. 2010) and is thought to function as a backup mechanism for trans-translation (Garza-Sanchez et al. 2011; Chadani et al. 2011; Schaub et al. 2012). Presence of ArfA in pathogenic bacteria, including *Salmonella, Yersinia* and *Klebsiella* species, underlines the importance of the ArfA-mediated ribosome rescue in stress adaptation and pathogenicity (Starosta et al. 2014). ArfA, a small protein with only 47 amino acids sufficient for function (Garza-Sanchez et al. 2011), recruits release factor RF2 to rescue stalled ribosomes (Chadani et al. 2012; Shimizu 2012). RF2 normally mediates translation termination at UGA or UAA stop codons by binding the stop codon and catalyzing peptidyl-tRNA hydrolysis and release of the nascent peptide (Korostelev 2011; Craigen and Caskey 1987). RF2 has remarkable specificity toward stop codons (Freistroffer et al. 2000) and does not function alone on truncated mRNA (Chadani et al. 2012; Shimizu 2012). How ArfA and RF2 sense the stalled ribosome, and how ArfA aids RF2 to catalyze peptide release in the absence of a stop codon is unknown.

To better understand ribosome rescue by ArfA and RF2, we formed an *E. coli* 70S ribosome rescue complex with mRNA truncated after an AUG codon in the P site, tRNA^fMet^, ArfA and RF2, and captured images of complexes by electron cryo-microscopy (cryo-EM; see Methods). Unsupervised classification of a single cryo-EM data set (Figure S1) using FREALIGN (Grigorieff 2016) revealed two ribosome structures with both ArfA and RF2 bound in the A site (Structures I and II; Figure 1 and Figure S2 and Table S1). Both structures contain tRNAs in the P and E (exit) sites and adopt a non-rotated conformation (Yusupov et al. 2001; Selmer et al. 2006; Korostelev et al. 2006), similar to that in translation termination complexes (Korostelev et al. 2008; Laurberg et al. 2008). High-resolution maps (Figures 1E, 1F and Figures S3-S5) allowed *de novo* modeling of ArfA (Figures 1F and Figure S3) and detailed structure determination of RF2 (Figure 1F and Figure S4) in each ribosome structure. The molecular interactions and conformational rearrangements inferred from Structures I and II provide the structural basis for ArfA-mediated ribosome rescue, as described below.

**Figure 1.**
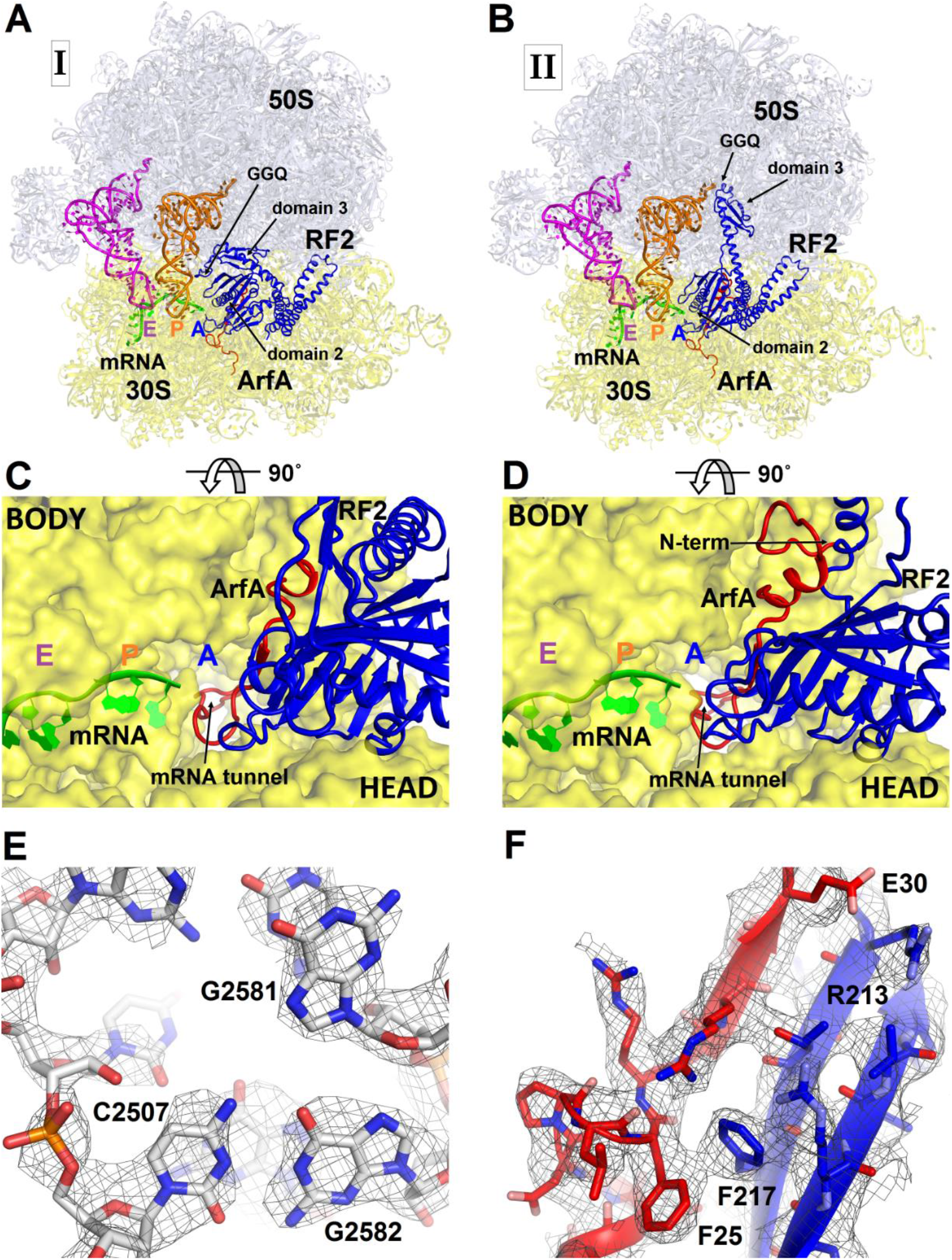
3.2-Å resolution cryo-EM structures of *E. coli* 70S ribosome bound with ArfA and release factor RF2. **(A)** Structure I with RF2 in a compact conformation; **(B)** Structure II with RF2 in an extended conformation. Domains 2 and 3 and the GGQ motif of RF2 are labeled. **(C)** and **(D)** A close-up view down the mRNA tunnel, showing RF2 and ArfA in the A site of Structure I **(C)** and Structure II **(D)**. The body and head domains of the 30S subunit are indicated. **(E)** Example of ribosomal RNA density (gray mesh) at the peptidyl-transferase center in Structure I, shown at σ=4.5 (23S rRNA nucleotides are labeled). (**F**) Extended β-sheet formed by ArfA (red model) and RF2 (blue model). Cryo-EM map (gray mesh) is shown for Structure II at σ=2.5. In all panels, the large 50S ribosomal subunit is shown in light blue; the small 30S subunit in yellow; mRNA in green; E-site tRNA in magenta; P-site tRNA in orange; ArfA in red and RF2 in blue. The maps were B-factor sharpened by applying the B-factor of -120 Å^2^. Additional views of cryo-EM density are available in figures S2, S3, S4 and S5.

## Results and Discussion

### ArfA C-terminus occupies the mRNA tunnel to sense the stalled ribosome

Sequence alignment of several hundreds of bacterial ArfA homologs reveals conserved hydrophobic N-terminal and positively charged C-terminal regions (Figure 2). In Structures I and II, the conformations of the N-terminal region differ as described in the following sections, but the rest of ArfA is similar. The mid-region of ArfA (His21 to Glu30; *E. coli* numbering is used) lies in the A site (Figures 1C-D). ArfA leaves a ~12-Å gap in the codon-binding region, sufficient to accommodate one or two nucleotides of mRNA following the P-site codon but not a longer mRNA, consistent with the reduced efficiency of ArfA-mediated release on mRNAs that extend three or more nucleotides beyond the P site (Zeng and Jin 2016).

**Figure 2.**
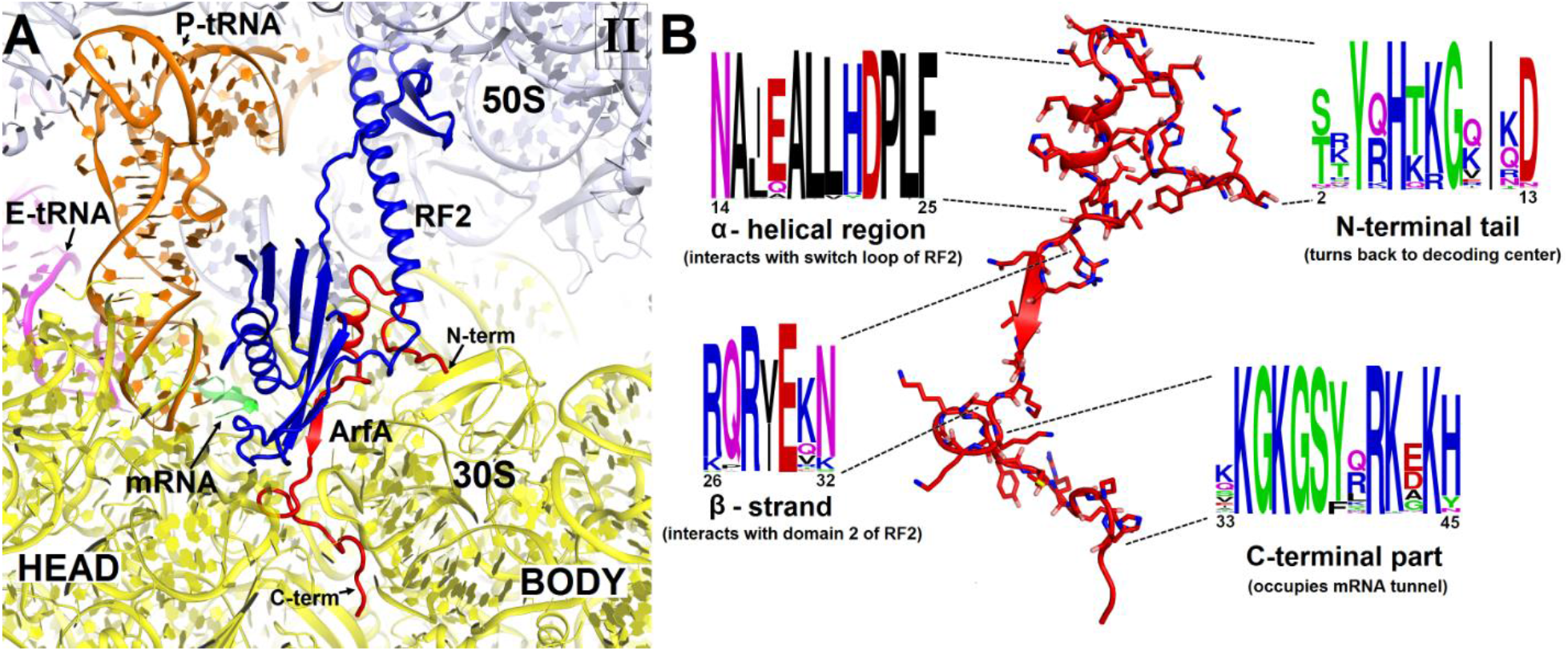
Structure and sequence of ArfA. **(A)** Close-up view of the intersubunit space and mRNA tunnel occupied by ArfA in Structure II. The color-coding is the same as in Fig.1. The head and body domains of the 30S subunit are indicated. (**B**) Sequence and structure of ArfA, shown in the same orientation as in (A). Sequence conservation among ~400 non-redundant bacterial ArfA homologs is shown for four ArfA regions (see Methods). The color code for amino acid type in the sequence logo is the following: green – polar, purple – neutral, blue – basic, red – acidic and black – hydrophobic residues.

The C-terminal region of ArfA occupies the mRNA tunnel between the head and body of the 30S subunit (Figures S6A and S7), as suggested by previous hydroxyl-radical probing studies (Kurita et al. 2014). The mRNA tunnel is primarily formed by the negatively charged 16S rRNA backbone. ArfA is stabilized by electrostatic interactions, as positively charged amino acids comprise nearly half of the ArfA residues within the mRNA tunnel (aa 31-48). ArfA residues Lys44 to Arg48 approach the mRNA tunnel entry at the solvent side of the 30S subunit, formed by 16S rRNA and proteins S3, S4, and S5 (Figure 2 and Figure S6A). Consistent with ArfA structure prediction (Kim, Chivian, and Baker 2004; Yang et al. 2015), the tail of ArfA appears to form an α-helix (aa ~50-55) next to S5, however the resolution of the map is not sufficient to build an unambiguous structural model (Figure S7). This suggests conformational disorder of the C-terminus at the entrance to the mRNA tunnel. Our structures are consistent with biochemical studies (Chadani et al. 2011), which showed that ArfA is functional with a C-terminal truncation at Asn47, but further shortening inactivates ArfA. In particular, truncations following Met40—removing at least five basic amino acids that bind in the tunnel—abrogate ArfA-mediated release by reducing ArfA affinity for the 70S ribosome (Chadani et al. 2011).

ArfA somewhat resembles the proteins ArfB (Gagnon et al. 2012) and SmpB (Ramrath et al. 2012; Neubauer et al. 2012), which mediate alternative rescue pathways (reviewed in (Keiler 2015)). In ArfB-mediated release and SmpB-mediated trans-translation, the proteins sense the stalled ribosomes by occupying the mRNA tunnel (Gagnon et al. 2012; Neubauer et al. 2012; Ramrath et al. 2012). Consistent with sequence divergence among the C-termini of ArfA, ArfB, and SmpB, however, each interacts with the mRNA tunnel differently (Gagnon et al. 2012; Neubauer et al. 2012; Ramrath et al. 2012), revealing how they mediate distinct ribosome rescue pathways (Figure S6).

### ArfA N-terminus is disordered in the presence of a compact (inactive) RF2 conformation

In Structure I, only the central and C-terminal parts of ArfA are visible, indicating that the N-terminal region (aa 2-16) is disordered. RF2 adopts a compact conformation (Figure 3A) similar to that of free RF2 (Figure 3B) (Vestergaard et al. 2001; Zoldak et al. 2007). By contrast, in canonical termination complexes formed on stop codons, release factors have only been observed in an extended (open) conformation (Korostelev 2011; Korostelev et al. 2008; Weixlbaumer et al. 2008). During translation termination, codon-recognition determinants in domain 2 (including the conserved ^205^SPF^207^ motif) of RF2 bind the stop codon in the A site of the 30S subunit. Helix α7 of domain 3 bridges the ribosomal subunits, placing the catalytic GGQ motif of domain 3 within the peptidyl-transferase center of the 50S subunit (Korostelev et al. 2008; Weixlbaumer et al. 2008). In the ArfA-bound Structure I, however, helix α7 packs on the β-sheet of domain 2 near the 30S subunit (Figures 3A and 3B). In this compact conformation, the loop that contains the ^250^GGQ^252^ motif of RF2 lies to the side of the β-sheet (near aa 165-168) of domain 2, facing the anticodon-stem loop and the D stem of the P-site tRNA. As such, the GGQ motif is roughly 70 Å away from its position within the peptidyl-transferase center in the canonical termination complex. Poor resolution of the catalytic GGQ residues (Figure S4A) at the tip of the loop suggests local structural flexibility, similar to that seen in crystal structures of free release factors (Shin et al. 2004; Vestergaard et al. 2001; Zoldak et al. 2007).

**Figure 3.**
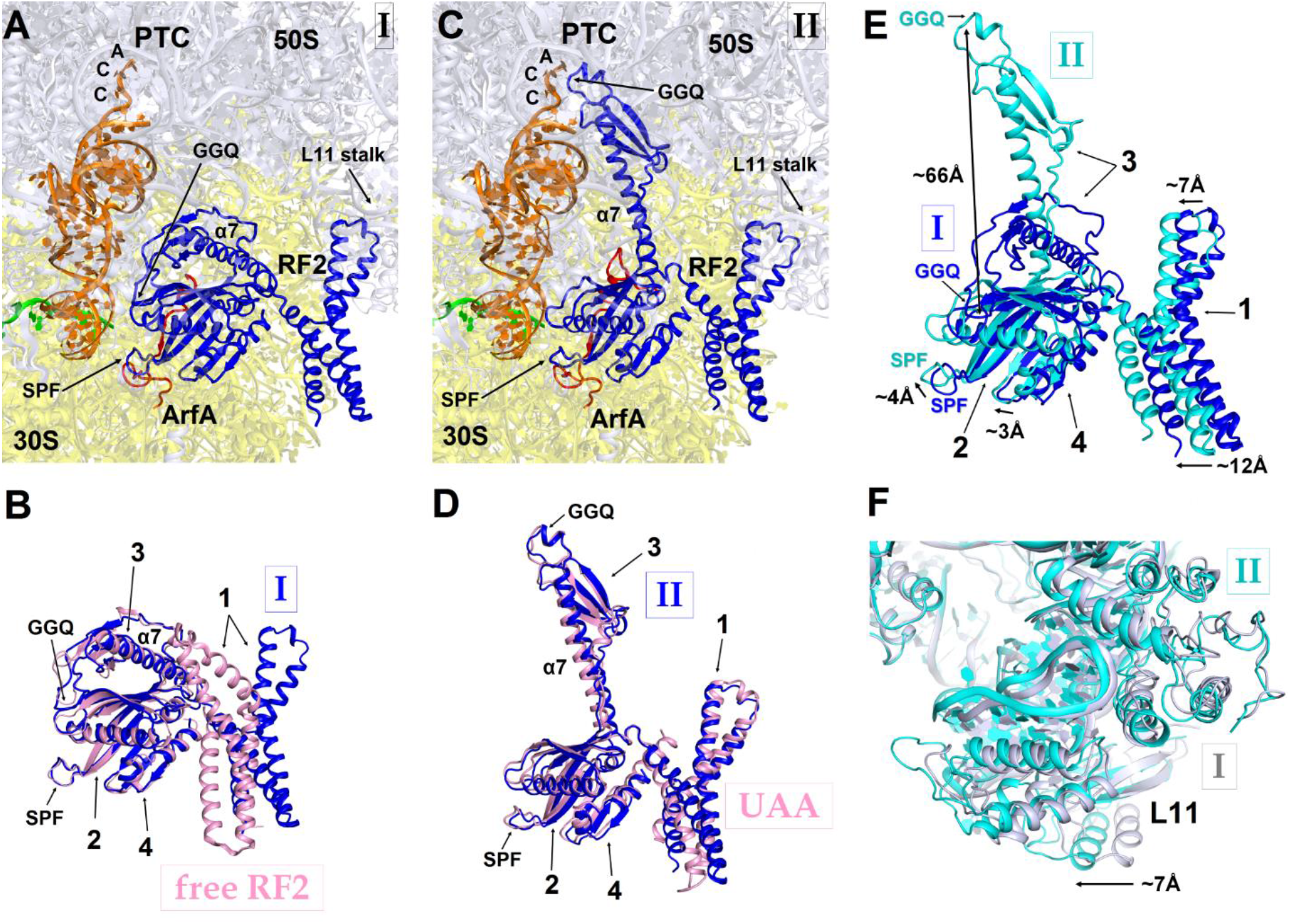
RF2 adopts two distinct conformations in Structures I and II. **(A)** The P and A sites of Structure I. ArfA is shown in red; RF2 in blue; mRNA in green; P-tRNA in orange; 30S subunit in yellow; and 50S subunit in light blue. **(B)** Superposition of RF2 from Structure I (blue) with the crystal structure of free (ribosome-unbound) *E. coli* RF2 (PDB 1GQE) (pink). Relative positions of the codon-recognition superdomain (domains 2 and 4) and catalytic domain 3 are nearly identical. The positions of domain 1 differ; this domain in both Structures I and II interacts with the L11 stalk at the 50S subunit shown in panels (A), (C) and (F). **(C)** The P and A sites of Structure II. The color coding is as in panel (A). **(D)** Superposition of extended RF2 in Structure II (blue) with *T. thermophilus* RF2 in the canonical termination complex formed on the UAA stop codon (PDB 4V67) (pink). The superposition was performed by structural alignment of 16S ribosomal RNAs. RF2 adopts similar conformations but domains 2 and 3 are positioned slightly differently with respect to the 30S subunit in the rescue complex II and in the termination complex (see also Figure 3). **(E)** Superposition of RF2 in Structures I (blue) and Structure II (cyan), achieved by structural alignment of the 16S ribosomal RNAs. Conformations of RF2 and positions relative to the 30S subunit differ between Structures I and II, as RF2 in Structure II binds deeper in the A site; differences in positions of RF2 regions are labeled with arrows. **(F)** Different positions of the L11 stalk, which interacts with domain 1 of RF2, in Structures I (light blue) and II (cyan), suggesting movement of the stalk together with domain 1 (E) upon RF2 activation. The view is similar to that shown in panels A, C and E. In panels (B), (D) and (E), the Arabic numerals label the domains of RF2.

The codon-recognition domain 2 of RF2 is positioned differently from that in canonical termination complexes. The domain is withdrawn from the A site, such that the SPF motif and other codon-recognition residues lie ~5 Å away from their positions in termination complexes bound to a stop codon (Figure 4 and Figure S4A) (Korostelev et al. 2008; Weixlbaumer et al. 2008). This position results from the mid-region of ArfA being sandwiched between domain 2 of RF2 and the decoding center. Here, the backbone of ArfA residues 25-29 binds to RF2 residues 213-217 within the β-sheet of domain 2 (Figure 1F), forming an extended β-sheet platform (Figures 4A and 4D). The conformation of the decoding center in Structure I differs from that in canonical termination complexes. In 70S termination complexes, the decoding center interacts with the switch loop of RF2 (aa 315-323), which bridges the codon-recognition and catalytic domains (Korostelev et al. 2008; Weixlbaumer et al. 2008). A1492 bulges from helix 44 (h44) of 16S rRNA and stacks on conserved Trp319 of the switch loop (Figure 4C), stabilizing the extended conformation of RF2 on the ribosome (Figure 4F) (Korostelev et al. 2008; Weixlbaumer et al. 2008). In Structure I, however, A1492 and A1493 lie inside h44 and are sandwiched between Pro23 of ArfA and A1913 of helix 69 of 23S rRNA (Figure 4A). The RF2 switch loop (at Trp319) is placed >10 Å away from its position in the termination complex and does not contact the decoding-center nucleotides (Figure 4D and Figure S4A).

**Figure 4.**
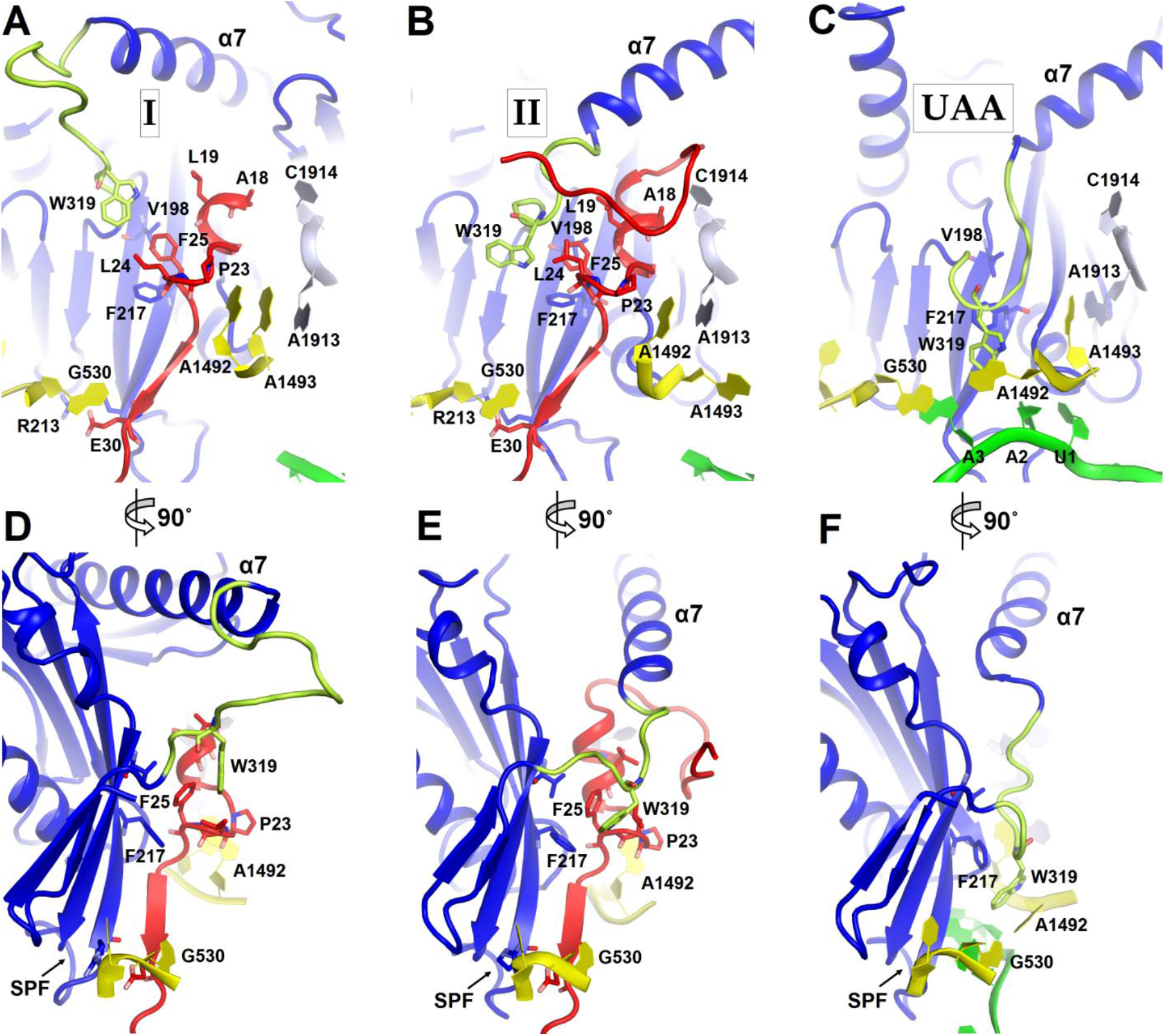
Positions of the codon-recognition domain (blue) and switch loop (yellow-green) of RF2 in Structures I and II (this work) and in the translation termination complex formed on the UAA stop codon (Korostelev et al. 2008). **(A-C)** Detailed view of the decoding center of Structure I (A), Structure II (B), and canonical termination complex formed on the UAA stop codon (C). **(D-F)** 90-degree rotated views (relative to that shown in panels A-C) of the decoding center in Structure I (D), Structure II (E) and the UAA-containing termination complex (F). The switch loop of RF2, which carries the conserved Trp319, adopts different positions in these three structures. Key structural features and residues of ArfA, RF2, mRNA stop codon and the ribosomal decoding center are labeled. ArfA is shown in red; RF2 in blue (RF2 switch loop in yellow-green); mRNA in green; 30S nucleotides in yellow; and 50S nucleotides in light blue.

Taken together, Structure I describes a 70S rescue complex in which ArfA stabilizes a compact form of RF2 resembling that of free RF2. The remote position of RF2’s catalytic GGQ motif—away from the peptidyl-transferase center—indicates that Structure I represents an inactive form of the 70S rescue complex.

### The ordering of the ArfA N-terminus is coupled with an extended (active) RF2 conformation

Structure II features an extended conformation of RF2 stabilized by interactions with the ordered N-terminal region of ArfA (Figures 1-4). The N-terminus of ArfA forms a minidomain, which packs between helix 69 of 23S rRNA, the β-sheet of domain 2, and extended helix α7 of RF2 (Figure S8). The tip of the ArfA minidomain (at aa 12-13) protrudes from the decoding center toward the 50S subunit, whereas the N-terminus folds back toward the decoding center, consistent with recent hydroxyl-radical probing studies (Kurita et al. 2014). Here, Thr7 and Lys8 bind h44 (at A1410) of 16S rRNA and the N-terminal amino acids bind the β-sheet of S12 (Figure S8).

A hydrophobic patch in the N-terminal minidomain of ArfA — formed by Leu19, Leu24, and Phe25—binds RF2 at Trp319 of the rearranged switch loop (Figures 4B, 4E and Figure S4B). These interactions explain the strict dependence of ArfA on RF2, rather than on the second release factor RF1 (Chadani et al. 2012; Shimizu 2012), whose switch loop is diverged from that of RF2 and lacks tryptophan (Korostelev et al. 2010). The hydrophobic patch in the N-terminal minidomain also binds RF2 at Val198 and Phe217 of the β-sheet of the codon-recognition domain 2 (Figures 4B, 4E and Figure S4B). In this configuration, the codon-recognition domain of RF2 partially settles into the decoding center, but remains ~3 Å from its position in canonical termination complexes bound to a stop codon. As in Structure I, the SPF motif remains unbound to the ribosome or ArfA. This observation explains why mutation of SPF motif residues—critical for stop-codon recognition—do not disrupt ArfA-mediated peptide release (Chadani et al. 2012).

The position of the ArfA N-terminal minidomain between RF2’s domain 2 and helix α7 of domain 3 results in docking of domain 3 into the peptidyl-transferase center (Figures 1, 3C, 3D). The opening of domain 3 is accompanied by movement of domain 1 of RF2 (Figure 3E), which binds the L11 stalk and shifts the L11 stalk by ~7 Å relative to that in Structure I (Figure 3F). The catalytic GGQ motif of domain 3 binds in the peptidyl-transferase center with the catalytic backbone amide of Gln252 (Korostelev et al. 2008) proximal to the ribose of the terminal A76 of P-site tRNA (Figure S4B). The conformation of the peptidyl-transferase center is nearly identical to that seen in canonical translation termination complexes (Korostelev et al. 2008). Structure II therefore represents an activated state of the ArfA•RF2-bound ribosome rescue complex.

Structures I and II are in agreement with previous biochemical and mutagenesis studies, as we have described above (Chadani et al. 2012; Chadani et al. 2010; Kurita et al. 2014; Zeng and Jin 2016). A single ArfA-inactivating mutation has been identified (Chadani et al. 2010). Mutation of Ala18 of the N-terminal ArfA minidomain to threonine prevents ArfA-mediated peptide release without disrupting RF2 binding (Shimizu 2012). In Structure II, Ala18 lies in the hydrophobic core of the N-terminal fold, tightly packed between the nucleobase of C1914 of h69 and Ile11 of ArfA (Figure S8). The substitution to the larger threonine residue is likely incompatible with the ordered N-terminal fold of ArfA and the extended conformation of RF2. The mutation should, however, be compatible with an inactive ribosome rescue complex (Structure I), consistent with RF2 binding and the lack of catalytic activity.

### Structural mechanism of ribosome rescue by ArfA and RF2

During ribosome rescue, the release of the nascent peptide should strictly coordinate with the recognition of a vacant mRNA tunnel. Our cryo-EM analysis indicates that ribosomes formed on a truncated mRNA in the presence of ArfA and RF2 interconvert between at least three equilibrium states, including Structures I, II and the ribosome with a vacant A site (Figure S1). These states suggest a structure-based model for step-wise release of nascent peptides from stalled ribosomes during ArfA-mediated ribosome rescue (Figure 5 and short movie also available at http://labs.umassmed.edu/korostelevlab/movarfa.gif).

**Figure 5.**
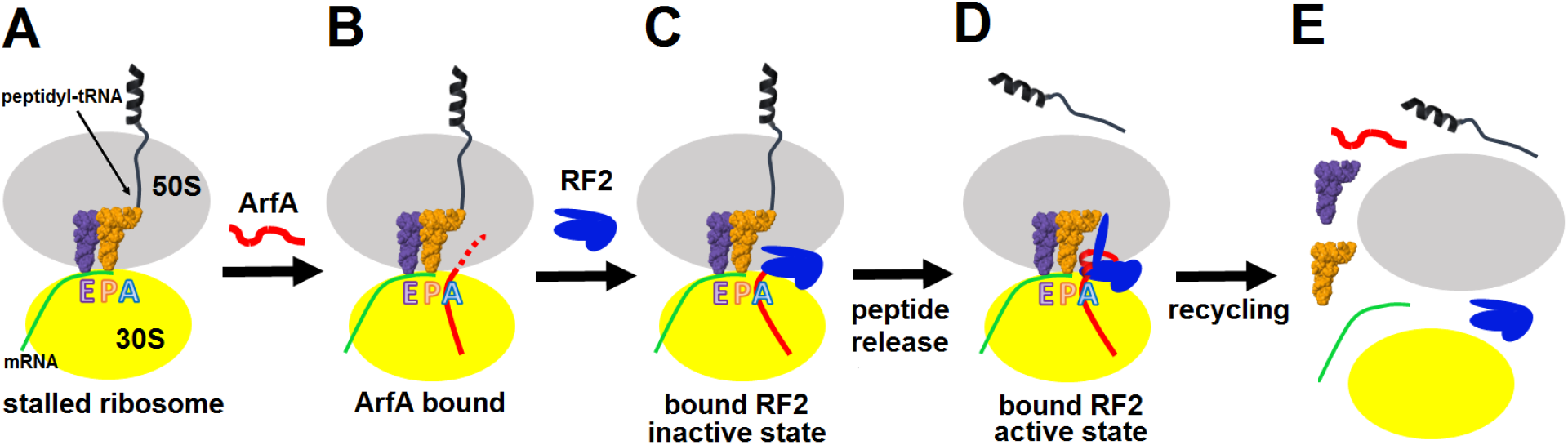
Mechanism of ArfA-mediated rescue of ribosomes stalled on truncated mRNA. **(A-B)** ArfA senses the stalled ribosome by binding its C-terminal portion in the vacant mRNA tunnel. **(C)** RF2 binds to the ribosome in inactive form (as in Structure I). **(D)** Folding of the N-terminal minidomain of ArfA is coupled with opening of RF2, so that the GGQ motif is placed into the peptidyl-transferase center (as in Structure II) and catalyzes peptidyl-tRNA hydrolysis, followed by **(E)** Recycling of the ribosome, likely mediated by the ribosome recycling factor.

In the absence of an A-site codon, peptidyl-tRNA-bound ribosomes cannot bind aminoacyl-tRNA and are recognized by ArfA, which binds the empty mRNA tunnel and recruits a compact (inactive) conformation of RF2 (Figure 5A-C). Biochemical studies have shown that ArfA can bind the ribosome without RF2 (Kurita et al. 2014), suggesting that ArfA precedes RF2 (Figure 5B-C). The initial rescue complex (Structure I) then samples an extended (active) conformation of RF2 coupled to ordering of the N-terminal minidomain of ArfA at the ribosomal decoding center (Structure II; Figure 5D). This structural rearrangement places the GGQ motif into the peptidyl-transferase center, as seen in canonical termination complexes (Korostelev et al. 2008; Weixlbaumer et al. 2008), where it catalyzes peptidyl-tRNA hydrolysis, releasing the nascent peptide from the ribosome (Figure 5D). Dissociation of ArfA and RF2, possibly through the inactive Structure I state, results in ribosomes with a vacant A site and deacylated tRNA in the P site, which enables recycling of the ribosome (Figure 5E).

The mechanism of ArfA-mediated ribosome rescue is remarkably different from canonical translation termination, wherein RF2 accurately defines the lengths of cellular proteins by direct recognition of stop codons in the A site (Korostelev et al. 2008; Weixlbaumer et al. 2008). ArfA converts RF2 into a stop-codon-independent release factor, but our structures show that: (i) ArfA does not mimic a stop codon; (ii) the conserved codon-recognition elements of RF2—including the SPF motif—are not required (Chadani et al. 2012); and (iii) instead of interacting directly with RF2, the ribosomal decoding center stabilizes ArfA, which in turn stabilizes an active RF2 conformation. Thus, bacteria have evolved an intricate stress-response mechanism in which a small protein with specific affinity to the stalled ribosome re-purposes a release factor. The ArfA-mediated ribosome rescue highlights an impressive ability of living organisms to co-opt the existing cellular mechanisms for different and sometimes mutually exclusive purposes.

## Materials and Methods

### Preparation of ArfA and RF2

The gene encoding *E. coli* ArfA (ASKA Clone(-) library, National BioResource Project, NIG, Japan) was subcloned into pET24b+ (Novagen) kanamycin resistance vector using the primer set CCCG<u>CATATG</u>CATCACCATCACCATCACATGAGTCGATATCAGCATACTAAAGGGC/CCCG<u>GGATCC</u>GTGATTTACTTTCTTGCCAC containing the NdeI/BamHI restriction sites (underlined) and transformed into an *E. coli* BLR/DE3 strain. The resulting ArfA protein is 60 amino acids long and is N-terminally His_6_-tagged. The cells containing the pET24b+ plasmid were cultured in Luria-Bertani (LB) medium with 50 µg mL^-1^ kanamycin at 37 °C until the OD_600_ reached 0.7-0.8. Expression of ArfA was induced by 1 mM IPTG (Gold Biotechnology Inc., USA), followed by cell growth for 9 hrs at 16 °C. The cells were harvested, washed and resuspended in buffer A (50 mM Tris-HCl (pH 7.5), 150 mM KCl, 10 mM imidazole, 6 mM β-mercaptoethanol (βME) and protease inhibitor (complete Mini, EDTA-free protease inhibitor tablets, Sigma Aldrich, USA). The cells were disrupted with a microfluidizer (Microfluidics, USA), and the soluble fraction was collected by centrifugation at 18000 rpm for 20 minutes and filtered through a 0.22 µm pore size sterile filter (CELLTREAT Scientific Products, USA).

ArfA was purified in three steps. The purity of the protein after each step was verified by 12 % SDS-PAGE stained with Coomassie Brilliant Blue R 250 (Sigma-Aldrich). First, affinity chromatography with Ni-NTA column (Nickel-nitrilotriacetic acid, 5 ml HisTrap, GE Healthcare) was performed using FPLC (Äkta explorer, GE Healthcare). The cytoplasmic fraction was loaded onto the column equilibrated with buffer A and washed with the same buffer. ArfA was eluted with a linear gradient of buffer B (buffer A with 0.5 M imidazole). Fractions containing ArfA were pooled and dialyzed against buffer C (buffer A without imidazole). The protein then was purified by ion-exchange chromatography (5ml HiTrap FF Q-column, GE Healthcare; FPLC). The column was equilibrated and washed with Buffer C, the protein was loaded in Buffer C and eluted with linear gradient of Buffer D (Buffer C with 1 M KCl). Finally, the protein was dialyzed against 50mM Tris-HCl buffer (pH 7.5), 150 mM KCl, 6 mM βME and protease inhibitor (complete Mini, EDTA-free protease inhibitor tablets, Sigma Aldrich, USA) and purified using size-exclusion chromatography (Hiload 16/60 Superdex 75pg column, GE Healthcare). The fractions of the protein were pulled, buffer exchanged (25 mM Tris-HCl (pH 7.0), 50 mM K(CH_3_COO), 10 mM Mg(CH_3_COO)_2_, 10 mM NH_4_(CH_3_COO) and 6 mM βME) and concentrated with an ultrafiltration unit using a 3-kDa cutoff membrane (Millipore). The concentrated protein was flash-frozen in liquid nitrogen and stored at -80 °C.

N-terminally His_6_-tagged RF2 (*E. coli* K12 strain) was purified as described (Korostelev et al. 2008; Laurberg et al. 2008).

### Preparation of the 70S rescue complex bound with ArfA•RF2

70S ribosomes were prepared from *E. coli* (MRE600) as described (Moazed and Noller 1986, 1989), and stored in the ribosome-storage buffer (20 mM Tris-HCl (pH 7.0), 100 mM NH_4_Cl, 12.5 mM MgCl_2_, 0.5 mM EDTA, 6 mM βME) at -80°C. Ribosomal 30S and 50S subunits were purified using sucrose gradient (10-35%) in a ribosome-dissociation buffer (20 mM Tris-HCl (pH 7.0), 300 mM NH_4_Cl, 1.5 mM MgCl_2_, 0.5mM EDTA, 6 mM βME). The fractions containing 30S and 50S subunits were collected separately, concentrated and stored in the ribosome-storage buffer at -80°C. *E. coli* tRNA^fMet^ was purchased from Chemical Block. RNA, containing the Shine-Dalgarno sequence and a linker to place the AUG codon in the P site (GGC AAG GAG GUA AAA <u>AUG</u>) was synthesized by IDT.

The 70S•mRNA•tRNA^fMet^•ArfA•RF2 complex was prepared by reconstitution *in vitro*. 2 µM 30S subunit (all concentrations are specified for the final solution) were pre-activated at 42°C for 5 minutes in the ribosome-reconstitution buffer (20 mM Tris-HCl (pH 7.0), 100 mM NH_4_Cl, 20 mM MgCl_2_, 0.5 mM EDTA, 6 mM βME). After pre-activation, 1.8 µM 50S subunit with 24 µM mRNA and 12 µM tRNA^fMet^ were added to the 30S solution and incubated for 15 minutes at 37°C. ArfA and RF2 were then added at 16 µM each and the solution was incubated for 15 minutes at 37°C and cooled down to room temperature. The solution was aliquoted, flash-frozen in liquid nitrogen and stored at -80 °C.

### Activity of ArfA and RF2

Activity of ArfA and RF2 in the ArfA-mediated rescue was tested using [^35^S]-formylmethionine release assay, essentially as we described previously (Svidritskiy et al. 2013). The pre-termination complex was formed as described (Svidritskiy et al. 2013), using *E. coli* 70S ribosomes, [^35^S]-labeled fMet-tRNA^fMet^ (^35^S-labeled methionine from Perkin Elmer) and truncated mRNA described above. Consistent with published data (Chadani et al. 2012; Shimizu 2012), neither ArfA nor RF2 alone induced release of [S^35^]-fMet from the pre-termination complex, whereas complete release was observed when ArfA and RF2 were added in combination.

### Cryo-EM and image processing

Holey-carbon grids (C-flat 2/2) were exposed to a 75% argon/25% oxygen plasma for 20 seconds using a Solarus 950 plasma cleaning system. The forward RF target was set to 7w. Before being applied to the grids, the 70S•mRNA•tRNA^fMet^•ArfA•RF2 complex was diluted in the ribosome-reconstitution buffer supplemented with ArfA and RF2 to the following final concentrations: ~0.45 µM 70S, 6 µM mRNA, 3 µM tRNA^fMet^, 10 µM ArfA and 10 µM RF2. 2 µl of the 70S•mRNA•tRNA^fMet^•ArfA•RF2 complex was applied to the grids. The grids were blotted for 5 s at blotting power 8 at 4°C and ~95% humidity and plunged into liquid ethane using an FEI Vitrobot MK4. The grids were stored in liquid nitrogen.

A dataset of 539,311 particles was collected as follows. 3760 movies were collected using Leginon (Suloway et al. 2005) on an FEI Krios microscope operating at 300 kV equipped with a DE-20 Camera System (Direct Electron, LP, San Diego, CA, USA) with -0.5 to -3.0 µm defocus. Each exposure was acquired with continuous frame streaming at 32 frames per second (fps) with various exposure lengths (38, 40, 54, 57 and 72 frames per movie) yielding a total dose of 61 e^-^/Å^2^. The nominal magnification was 29,000 and the calibrated pixel size at the specimen level was 1.215 Å. The frames for each movie were processed using DE_process_frames script (in EMAN2 (Tang et al. 2007)) which is available from Direct Electron at http://www.directelectron.com/scripts. The movies were motion-corrected and frame averages were calculated using the first half of each movie (data up to a dose of ~30 e^-^/Å^2^) and excluding the first two frames, after multiplication with the corresponding gain reference. CTFFIND4 (Rohou and Grigorieff 2015) was used to determine defocus values for each resulting frame average. 503 movies with large drift, low signal, heavy ice contamination, or very thin ice were excluded from further analysis after inspection of the averages and the power spectra computed by CTFFIND4. Particles were semi-automatically picked from full-sized images in EMAN2 using ~50 particles picked manually to serve as a reference. 320x320 pixel boxes with particles were extracted from images and normalized. The stack and FREALIGN parameter file were assembled in EMAN2. To speed up data processing, a 4x-binned image stack was prepared using EMAN2.

Data classification is summarized in Extended Data – Fig. 1. FREALIGN v9 (versions 9.10-9.11) was used for all steps of refinement and reconstruction (Grigorieff 2016). The 4x-binned image stack (539,311 particles) was initially aligned to a ribosome reference (PDB 4V4A) (Vila-Sanjurjo et al. 2003) using 3 cycles of mode 4 (search and extend) alignment including data in the resolution range from 300 Å to 30 Å until the convergence of the average score. Subsequently, the 4x binned stack was aligned against the common reference resulting from the previous step, using mode 1 (refine) in the resolution ranges 300-18 Å and 300-12 Å (for both ranges, 3 cycles of mode 1 were run). In the following steps, the 4x binned stack was replaced by the unbinned (full-resolution) image stack, which was successively aligned against the common reference using mode 1 (refine), including gradually increasing resolution limits (increments of 1 Å, 5 cycles per each resolution limit) up to 6 Å. The resolution of the resulting common reference was 3.29 Å (Fourier Shell Correlation (FSC) = 0.143). Subsequently, the refined parameters were used for classification of the unbinned stack into 10 classes in 30 cycles using the resolution range of 300-6 Å. This classification revealed 7 high-resolution classes and 3 low-resolution (junk) classes (Figure S1). The particles assigned to the high-resolution classes that contained RF2 and ArfA were extracted from the unbinned stack (with > 50% occupancy and scores > 0) using merge_classes.exe (part of the FREALIGN distribution), resulting in a stack containing 320,895 particles. Classification of this stack was performed for 30 cycles using a focused spherical mask around the A site (55 Å radius, as implemented in FREALIGN). This classification yielded 3 high-resolution classes, two of which contained both ArfA and RF2 (Structures I and II) and one with a vacant mRNA tunnel and A site. Using more classes (up to 8) did not yield additional structures (e.g. containing ArfA alone, RF2 alone or additional ArfA•RF2 conformations). For the classes of interest (Structures I and II), particles with > 50% occupancy and scores > 0 were extracted from the unbinned stack. Refinement to 6 Å resolution using mode 1 (5 cycles) resulted in ~3.15 Å maps (FSC=0.143). The maps were B-factor sharpened using automatically calculated B-factors (approximately -90 Å^2^) in bfactor.exe (part of the FREALIGN distribution) and used for model building and structure refinements. B-factors of -120 or -150 Å^2^ were also used to interpret high-resolution details in the ribosome core regions. FSC curves were calculated by FREALIGN for even and odd particle half-sets.

### Model building and refinement

Recently reported cryo-EM structure of *E. coli* 70S•RelA•A/R-tRNA^Phe^ complex (Loveland et al. 2016), excluding RelA and tRNA^Phe^, was used as a starting model for structure refinement. The structure of compact RF2 (Structure I) was built using the crystal structure of free RF2 (PDB 1GQE) (Vestergaard et al. 2001) as a starting model. The extended form of RF2 (Structure II) was created by homology modeling from *Thermus thermophilus* RF2 within a 70S termination complex (Korostelev et al. 2008) using SWISS-PROT (Bairoch et al. 2004). ArfA was modeled *de novo* in Coot (Emsley and Cowtan 2004), using an initial structure predicted by ROBETTA (Kim, Chivian, and Baker 2004). The secondary structure in our resulting model of AfrA is consistent with those predicted by ROBETTA and I-TASSER (Yang et al. 2015). Initial protein and ribosome domain fitting into cryo-EM maps was performed using Chimera (Pettersen et al. 2004), followed by manual modeling using Pymol (DeLano 2002) and Coot. The linkers between the domains and parts of the domains that were not well defined in the cryo-EM maps (e.g. loops of RF2 in Structure I, shown in Figure S4A) were modeled as protein or RNA backbone.

Structures I and II were refined by real-space simulated-annealing refinement using atomic electron scattering factors (Gonen et al. 2005) in RSRef (Chapman 1995; Korostelev, Bertram, and Chapman 2002) as described (Svidritskiy et al. 2014). Secondary-structure restraints, comprising hydrogen-bonding restraints for ribosomal proteins and base-pairing restraints for RNA molecules, were employed as described (Korostelev et al. 2008). Refinement parameters, such as the relative weighting of stereochemical restraints and experimental energy term, were optimized to produce the stereochemically optimal models that closely agree with the corresponding maps. In the final stage, the structures were refined using phenix.real_space_refine (Adams et al. 2010), followed by a round of refinement in RSRef applying harmonic restraints to preserve protein backbone geometry. Ions were modeled as Mg^2+^, filling the difference-map peaks using CNS (Brunger 2007). To this end, the maps were converted to structure factors using phenix.map_to_structure_factors (Adams et al. 2010). The refined structural models closely agree with the corresponding maps, as indicated by low real-space R-factors of 0.18 and 0.19 for Structures I and II, respectively. The resulting models have excellent stereochemical parameters (100^th^-percentile MolProbity score), characterized by low deviation from ideal bond lengths and angles, low number of protein-backbone outliers (no outliers in ArfA) and other robust structure-quality statistics, as shown in Table S1. Structure quality was validated using MolProbity (Chen et al. 2010).

Structure superpositions and distance calculations were performed in Pymol. Figures were prepared in Pymol and Chimera (DeLano 2002; Pettersen et al. 2004).

### Sequence and structural analysis

NCBI (PSI) BLAST (Altschul et al. 1997) was used to obtain ~400 non-redundant ArfA homolog sequences with less than 95% identity to that of *E. coli* ArfA. MUSCLE (Edgar 2004) was used to generate a multiple-sequence alignment which was presented with WebLogo 3 (Crooks et al. 2004) (Figure 2).

## Acknowledgments

We thank the National Institute of Genetics, Shizouka, Japan for providing ArfA- and RF2-overexpressing strains of *E. coli* (ASKA- library); Michael Spilman (Direct Electron, LP) for assistance with data collection and processing; Anna B. Loveland for assistance with data processing; Darryl Conte Jr. for assistance with manuscript preparation; members of the Korostelev laboratory for helpful discussions and comments on the manuscript. This study was supported by NIH Grants R01 GM106105 and GM107465 (to A.A.K.).

## Author Contributions

GD and AAK conceived and designed the project; GD prepared the ribosome complex and processed cryo-EM data; GD and AAK built structural models and wrote the manuscript; ES and RM assisted with protein and ribosome purification; ES and GD tested ArfA and RF2 in peptide release assays; RD, TG and NG assisted with cryo-EM data collection and processing of preliminary data sets; DS assisted with cryo-EM sample preparation, collected cryo-EM data and assisted with initial data processing; all authors contributed to manuscript finalization.

## Supplementary figures and tables

**Figure S1.**
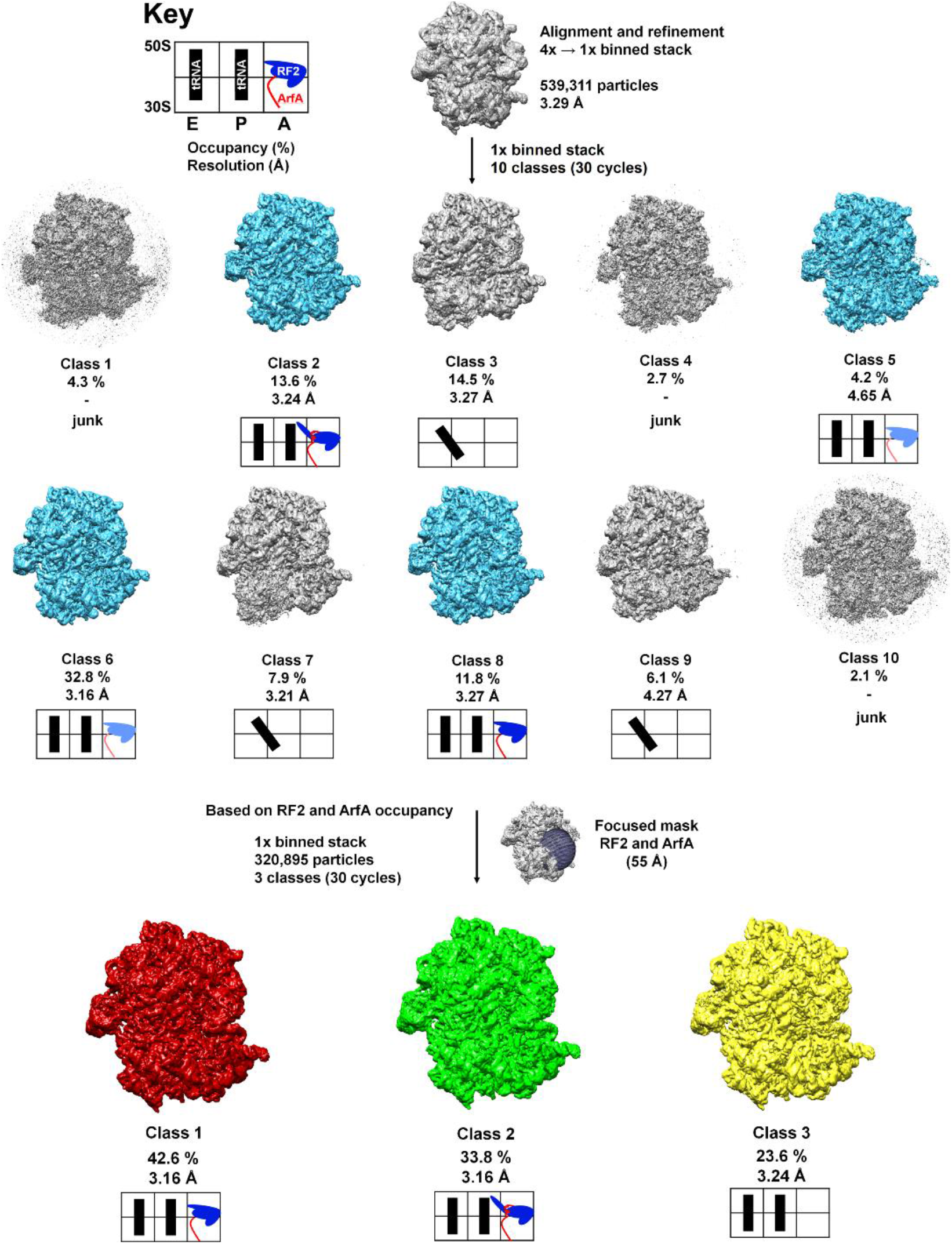
Schematic of cryo-EM refinement and classification. All particles (4x stack) were initially aligned to a single model. 3D classification (10 classes) using the unbinned (1x binned) stack was used to identify the particles containing ArfA and RF2. Subsequent 3D classification using a spherical mask around the A site yielded 3 “purified” classes representing Structure I, Structure II and the ribosome with a vacant A site. Additional sub-classifications (including up to eight classes) did not yield additional structures (e.g. ArfA-, RF2- or ArfA•RF2-containing classes). Light blue color for RF2 means partial occupancy of RF2.

**Figure S2.**
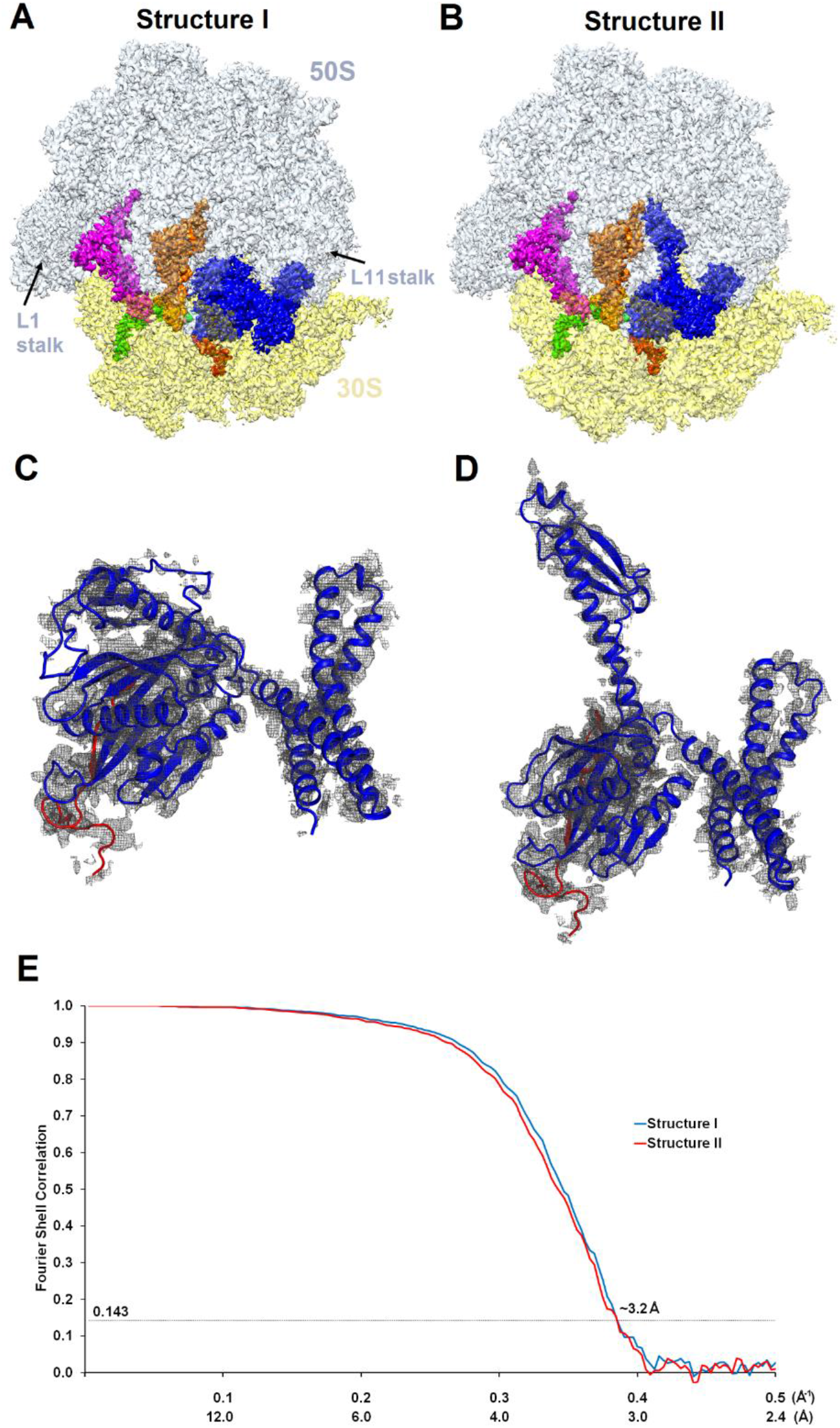
Cryo-EM densities of Structures I and II. (**A**) and (**B**) Cryo-EM maps of 70S•ArfA•RF2 complexes were segmented and colored as in Figure 1. (**C**) and (**D**) Cryo-EM density (gray) for ArfA (red model) and RF2 (blue model) in Structures I and II. The maps were B-factor sharpened by applying the B-factor of -120 Å^2^. (**E**) Fourier shell correlation as a function of resolution for Structures I and II.

**Figure S3.**
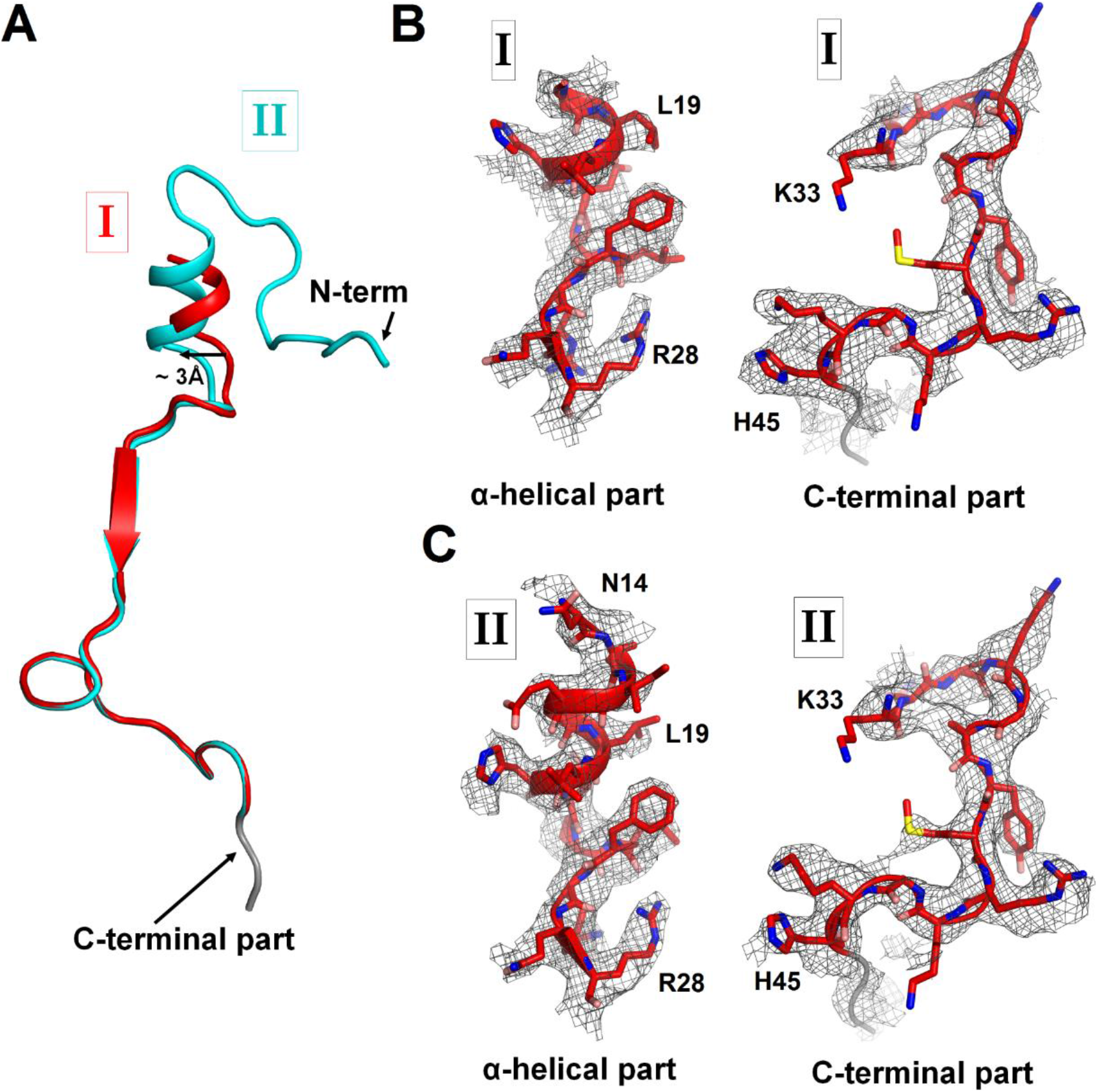
Cryo-EM densities corresponding to N- and C-terminal regions of ArfA in Structures I and II. (**A**) Superposition of ArfA in Structures I (red) and Structure II (cyan), achieved by structural alignment of the 16S ribosomal RNA. Conformations of the N-terminal region of ArfA relative to the 30S subunit differ between Structures I and II; the difference in the position of the α-helical part is shown with an arrow. (**B**) Partially resolved α-helical part at the N-terminus and the C-terminal part of ArfA in Structure I (red model) defined by cryo-EM map (gray mesh) at σ=2.5. (**C**) Well-resolved α-helical part at the N-terminus and the C-terminal part of ArfA in Structure II (red model) defined by cryo-EM map (gray mesh) at σ=2.5. The maps were B-factor sharpened by applying the B-factor of -120 Å^2^. A poorly defined region (with the backbone traceable at low σ=1.0) of the C-terminal structure is colored in gray.

**Figure S4.**
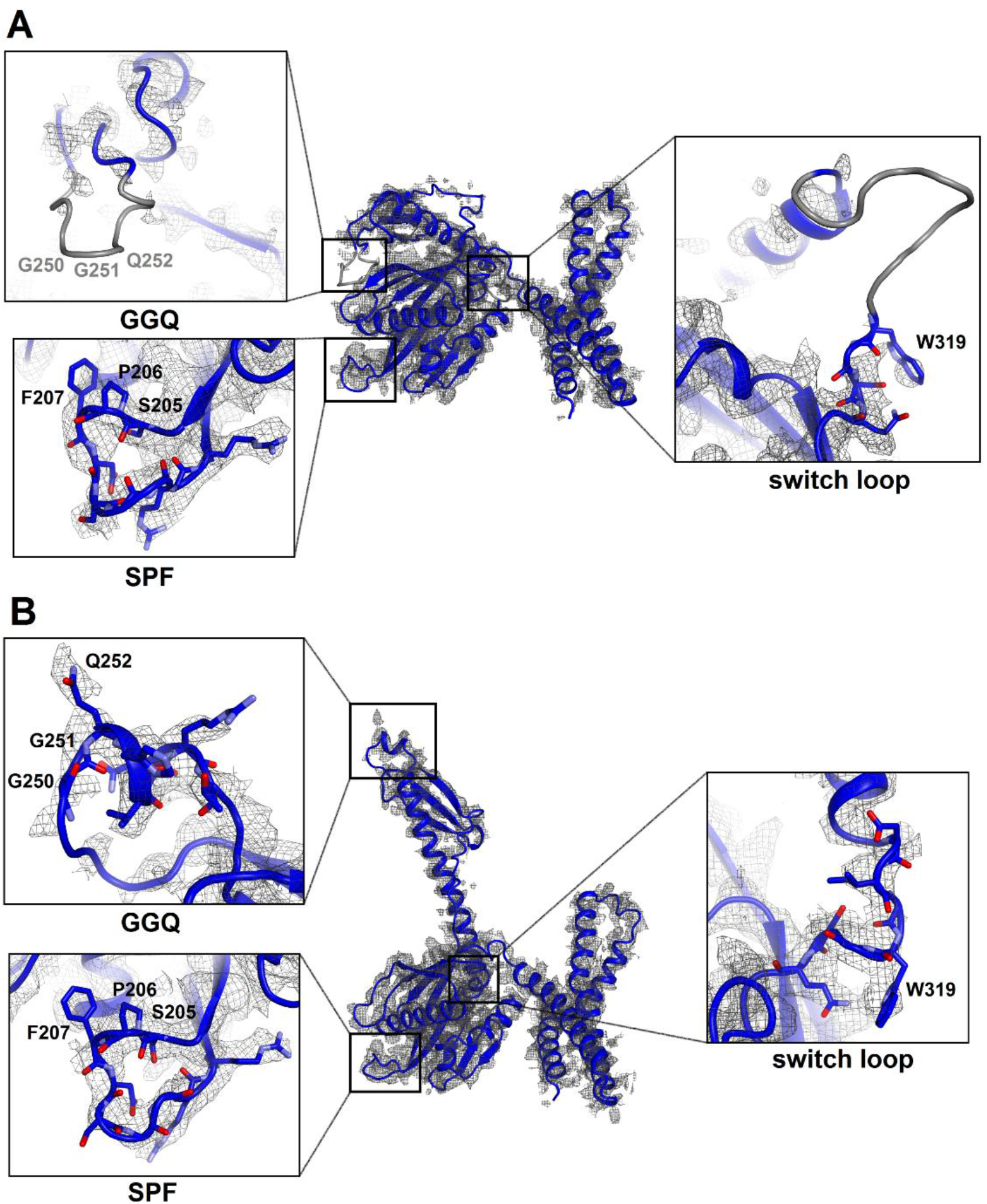
Cryo-EM densities corresponding to functional regions of RF2 in Structures I and II. (**A**) Compact conformation of RF2 (blue model) defined by cryo-EM map (gray mesh) at σ=2.5. Close-up views are shown in boxes for the SPF and GGQ motifs and the switch loop (the orientations differ from that in the main panel) Gray regions are poorly defined in the map, in that the backbone is only traceable at low σ=1.0 or lower. (**B**) Extended conformation of RF2 defined by cryo-EM map at σ=2.5. Close-up views are shown in boxes for the SPF and GGQ motifs and the switch loop. The maps were B-factor sharpened by applying the B-factor of -120 Å^2^.

**Figure S5.**
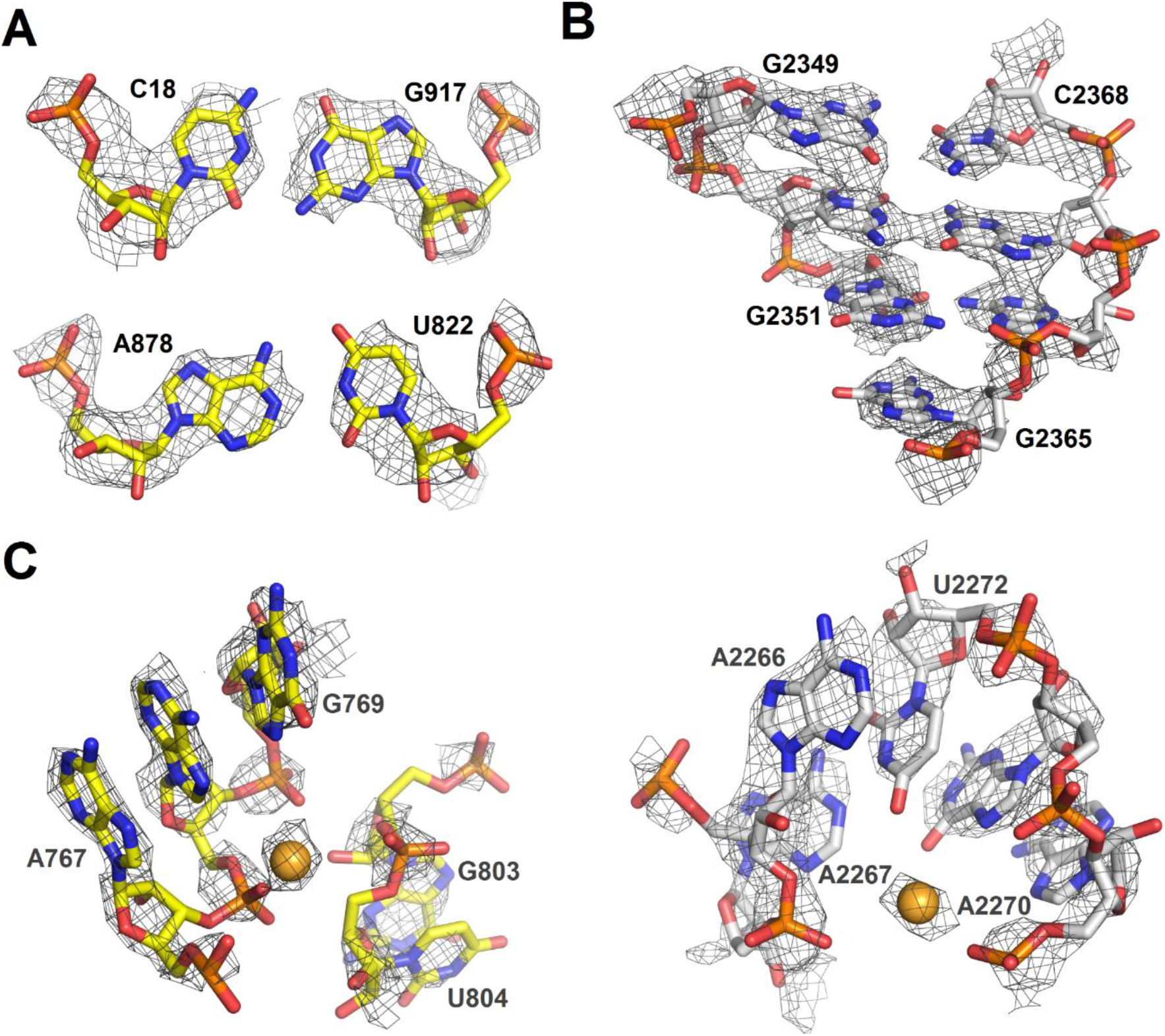
Cryo-EM densities of ribosomal RNA in Structure II. (**A**) Density (gray mesh) at σ=4.5 for 16S rRNA nucleotides forming base pairs. (**B**) Density (gray mesh) at σ=4.5 for 23S rRNA (part of helix 86) (**C**) Densities (gray mesh) for a magnesium ion at σ=4.5 (gold sphere) coordinated by 16S rRNA (yellow) at σ=6.0 or 23S rRNA (gray) at σ=6.0. The maps were B-factor sharpened by applying the B-factor of -150 Å^2^.

**Figure S6.**
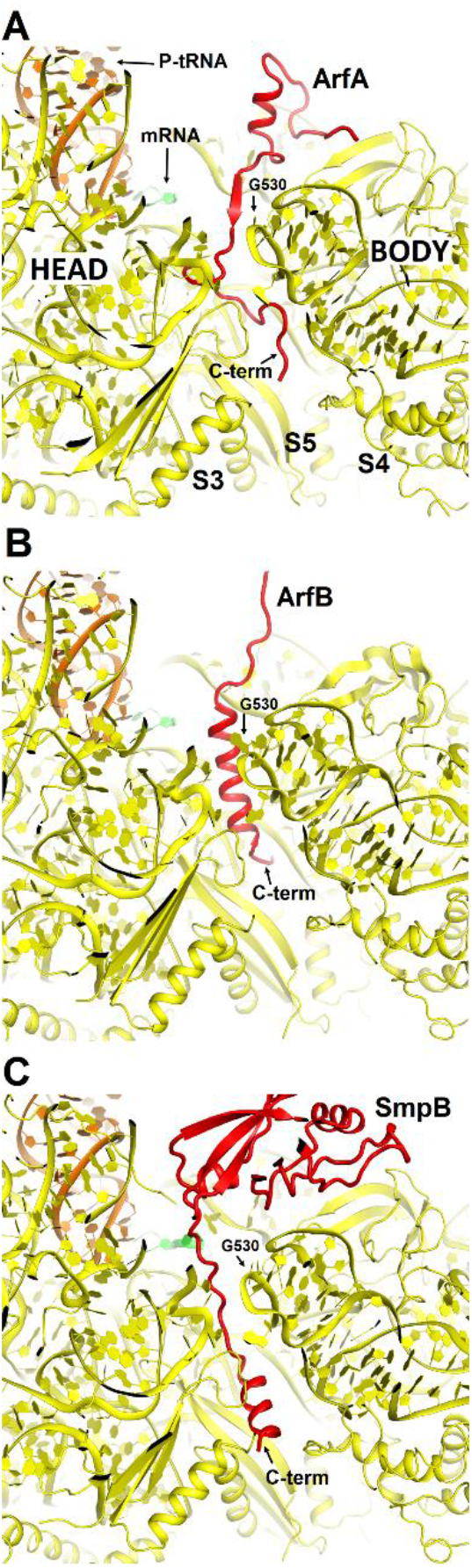
Comparison of mRNA tunnel occupancies by ribosome rescue proteins ArfA, ArfB and SmpB (shown in red). (**A**) Structure of ArfA and interactions with the 30S subunit in Structure II (this work), (**B**) Structure of ArfB and interactions with the 30S subunit in the crystal structure of the 70S-bound ArfB (PDB 4V95) (**C**) Structure of SmpB and interactions with the 30S subunit in the crystal structure of the 70S-bound SmpB and tmRNA mimic (PDB 4V8Q). In all panels, the 30S subunit is shown in yellow, P-site tRNA in orange and mRNA in green. The entrance to the mRNA tunnel (at the solvent interface of the 30S subunit) is formed by proteins S3, S4 and S5, which are labeled in panel A. G530 is labeled in each panel to show the location of the decoding center.

**Figure S7.**
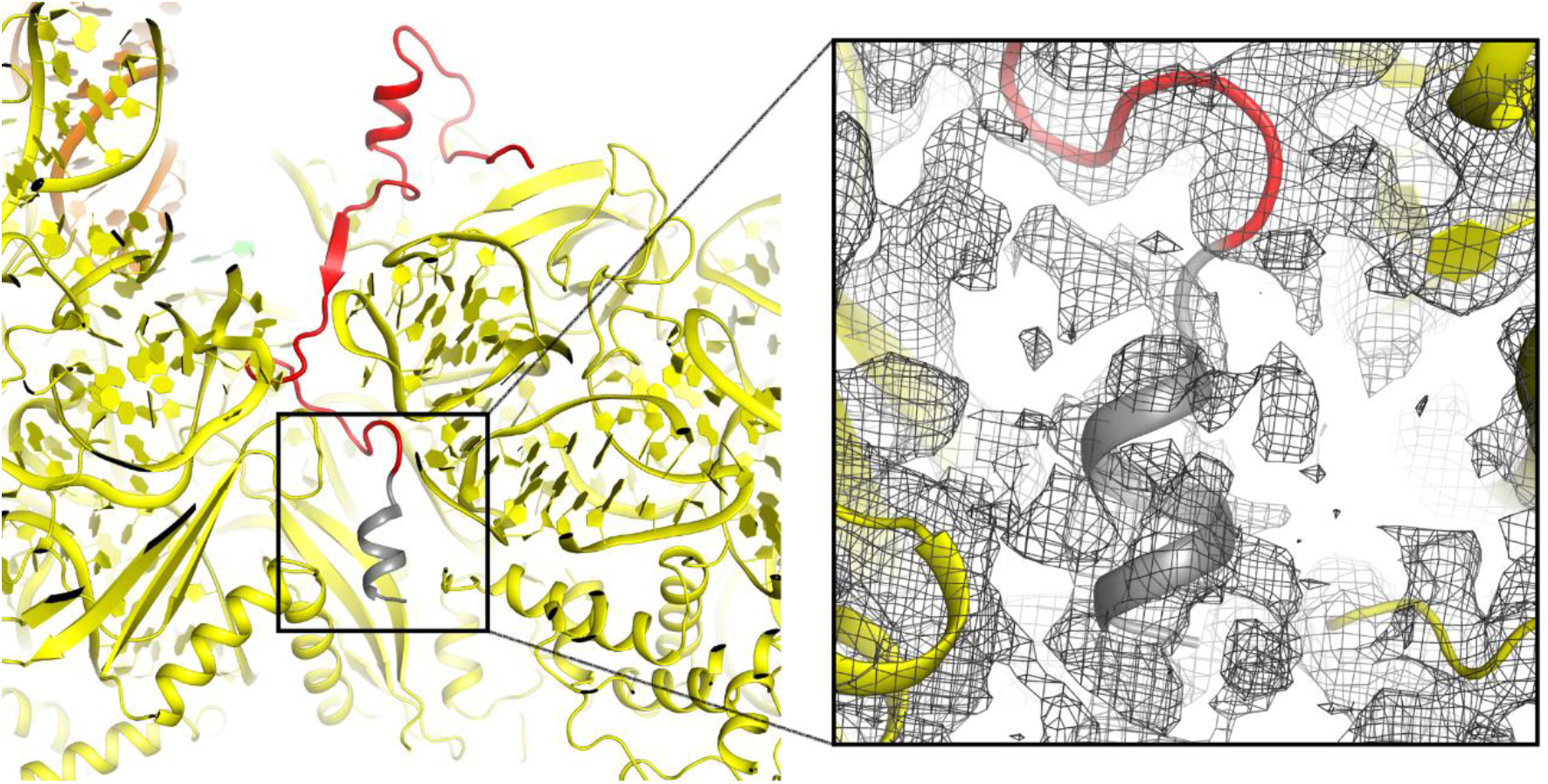
Putative α-helical structure of the C-terminal region of ArfA at the entry of the mRNA tunnel (near protein S5) in Structures I and II (Structure II is shown). The view is similar to that shown in Fig. S6A. The α-helix at the C-terminal tail of ArfA is predicted by ROBETTA and I-TASSER. The putative structural model (aa ~47-55) is colored in gray and defined by cryo-EM map (gray mesh) at σ=2.0. The map was B-factor sharpened by applying the B-factor of -120 Å^2^. The 30S subunit is shown in yellow, the well-ordered part of ArfA (up to aa 46) is in red.

**Figure S8.**
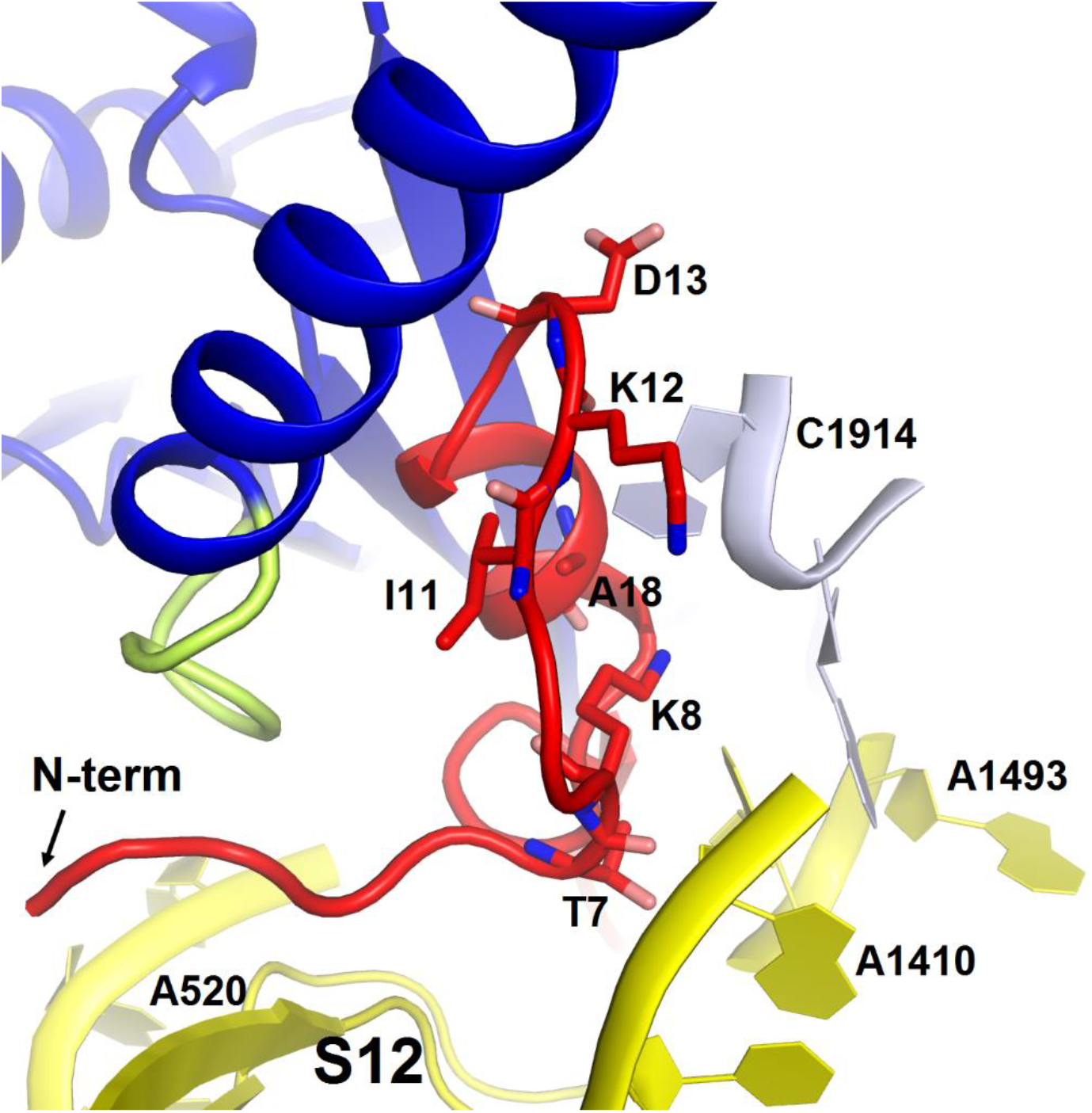
Structure of the N-terminal tail of ArfA near the decoding center in Structure II. The N-terminus of ArfA (red) is positioned next to A520 of helix 18 of 16S rRNA (within ~10 Å) and contacts S12. Polar residues Thr7 and Lys8 interact with h44 of 16S rRNA (at A1410) and h69 of 23S RNA (at C1914), respectively. The 30S subunit is shown in yellow, 50S in light blue, RF2 in blue. The switch loop of RF2 is highlighted in yellow-green.

**Table S1.** Cryo-EM data collection and refinement statistics

|  | Structure I | Structure II |
| --- | --- | --- |
| **Data collection** |  |  |
| EM equipment | FEI Titan Krios | FEI Titan Krios |
| Voltage (kV) | 300 | 300 |
| Detector | DE-20 | DE-20 |
| Pixel size (Å) | 1.215 | 1.215 |
| Electron dose (e<sup>-</sup>/Å<sup>2</sup>) | 61 (used 30) | 61 (used 30) |
| Defocus range (μm) | -0.5 to -3.0 | -0.5 to -3.0 |
| **Reconstruction** |  |  |
| Software | Frealign v9.10-9.11 | Frealign v9.10-9.11 |
| Number of particles used | 139,861 | 96,070 |
| Final resolution (Å) | 3.15 | 3.15 |
| Map-sharpening *B* factor (Å<sup>2</sup>) | -92.1 | -89.9 |
| **Model building** |  |  |
| Software | Coot | Coot |
| **Model composition** |  |  |
| Non-hydrogen atoms | 153057 | 153192 |
| Protein residues | 6556 | 6571 |
| RNA bases | 4727 | 4727 |
| Ligands (Zn<sup>2+</sup>/Mg<sup>2+</sup>) | 2/348 | 2/360 |
| **Refinement** |  |  |
| Software | RSRef and Phenix | RSRef and Phenix |
| Correlation Coeff.(%) | 81.98 | 80.34 |
| R-factor | 0.183 | 0.191 |
| **Validation (proteins)** |  |  |
| MolProbity score | 2.2 (100<sup>th</sup> percentile) | 2.2 (100<sup>th</sup> percentile) |
| Clash score, all atoms | 12.0 (97<sup>th</sup> percentile) | 12.2 (97<sup>th</sup> percentile) |
| Good rotamers (%) | 93.8 | 94.5 |
| Ramachandran-plot statistics (%) |  |  |
| Favored (overall) | 88.4 | 88.1 |
| Allowed (overall) | 10.6 | 10.8 |
| Outlier (overall) | 1.0 | 1.1 |
| Favored (ArfA) | 86.7 | 88.9 |
| Allowed (ArfA) | 13.3 | 11.1 |
| Outlier (ArfA) | - | - |
| Favored (RF2) | 86.1 | 92.5 |
| Allowed (RF2) | 13.9 | 7.2 |
| Outlier (RF2) | - | 0.3 |
| R.m.s. deviations |  |  |
| Bond length (Å) | 0.004 | 0.004 |
| Bond angle (°) | 0.842 | 0.864 |
| **Validation (RNA)** |  |  |
| Correct sugar puckers (%) | 99.8 | 99.7 |
| Good backbone conformation (%) | 85.1 | 85.0 |

